# Long-term associative memory in rats: effects of familiarization period in object-place-context recognition test

**DOI:** 10.1101/728295

**Authors:** Shota Shimoda, Takaaki Ozawa, Yukio Ichitani, Kazuo Yamada

## Abstract

Spontaneous recognition tests, which utilize rodents’ innate tendency to explore novelty, can evaluate not only simple non-associative recognition memory but also more complex associative memory in animals. In the present study, we investigated whether the length of the object familiarization period (sample phase) improved subsequent novelty discrimination in the spontaneous object, place, and object-place-context (OPC) recognition tests in rats. In the OPC test, rats showed a significant novelty preference only when the familiarization period was 30 min but not when it was 5 min or 15 min. However, the rats exhibited a successful discrimination between the stayed and replaced objects under 15 min and 30 min familiarization period conditions in the place recognition test and between the novel and familiar objects under all conditions of 5, 15 and 30 min in the object recognition test. Our results suggest that the extension of the familiarization period improves performance in the spontaneous recognition paradigms, and a longer familiarization period is necessary for long-term associative recognition memory than for non-associative memory.

## Introduction

Recognition memory is necessary to discriminate novel information from what is already known. Since animals have an innate tendency to respond to or explore novel stimuli, the habituation-dishabituation paradigm has been regarded as a useful behavioral test to assess recognition memory in various animal species including Aplysia (Pinsker, Kupfermann, Castellucci, and Kandel 1970), rodents (Ennaceur and Delacour 1988), monkeys (Mishkin and Delacour 1975), and humans (Fantz 1964). In particular, researchers have evaluated rodents’ recognition memory using several types of spontaneous recognition tests.

Spontaneous recognition tests have been used to evaluate not only simple non-associative recognition memory (e.g. object recognition test, Ennaceur and Delacour 1988; place recognition test, Ennaceur and Meliani 1992) but also more complex associative recognition memory (e.g. object-context recognition test, Dix and Aggleton, 1999; object-place-context (OPC) recognition test, Eacott and Norman 2004). While non-associative recognition memory is typically composed of single elements of information such as objects or locations, associative recognition memory is necessary to recognize objects using combined information on multiple elements that typically include the context where animals encountered the objects, as well as the information on the objects and locations.

A standard object recognition test consists of a sample phase and a test phase, with a delay period inserted between the two phases. In the sample phase, a rat is allowed to explore an open-field arena, in which a pair of two identical objects are placed, for a few minutes for familiarization. After the delay period, the rat is returned to the arena where one of the objects is replaced with a novel object (test phase). A preferential exploration toward the novel object is defined as a successful discrimination, and the rat is considered to exercise a great ability of recognition memory. In general, performance in the test depends on the length of the delay period, such that a shorter (< 24 hours) and a longer (≧24 hours) delay period reflects short-term and long-term recognition memory, respectively (Ennaceur and Delacour 1988). Moreover, performance in the spontaneous recognition memory test could be affected by the length of the familiarization period (sample phase). Previous studies systematically examined how the extension of familiarization periods improved performance in non-associative object or place recognition tests. For example, in an object recognition test, animals showed novel object preference in a 5 min familiarization with both short (15 min) and long (24 hours) delay periods (Gaskin et al. 2010). Likewise, in a place recognition test with a long (24 hours) delay period, a 20 min, but not 5 min familiarization was sufficient for rats to discriminate a replaced object from a stayed one (Ozawa et al. 2011).

In associative recognition memory tests, such as the OPC recognition test, however, rats would identify an association between objects, places, and contexts. Typically, the OPC recognition test consists of two sample phases, a test phase, and a delay inserted between the second sample and test phases. In the first sample phase, rats are familiarized with two different objects in a context. In the second sample phase, the rats are moved to another context in which the same pair of objects are placed in a swapped position. Subsequently, in the test phase, then, the rats are placed in one of the contexts and allowed to explore a pair of one of the objects that the rats encountered in either the first or the second contexts. In the OPC recognition test, rats are expected to explore the replaced object longer than the one that stayed in each context. Several studies demonstrated that short-term recognition memory had been evident in the OPC recognition test which consisted of 2-5 min familiarizations and 2-15 min delay periods (Davis et al. 2013; Eacott and Norman 2004; Easton et al. 2011; McLean et al. 2018; Ramsaran et al. 2016; Langston and Wood 2010; Cozannet et al. 2010; Lee et al. 2014; Wilson et al. 2013), although long-term recognition memory (≧24 hours) has never been tested.

In the present study, we hypothesized that (1) the extension of the familiarization period in the sample phase facilitated associative recognition memory and enabled animals to exhibit long-term associative recognition memory in the OPC recognition test, and (2) a longer familiarization period was necessary for the formation of long-term associative recognition memory than for the non-associative memory. Here, we systematically investigated the relationship between the lengths of familiarization periods (5, 15, or 30 min) and subsequent novelty discrimination performance in the object, place or OPC recognition tests in rats.

## Materials and methods

### Subjects

Thirty-two male Long-Evans rats (10-11 weeks old and weighing 346.05 ± 11.61 g at the beginning of behavioral experiments (Institute for Animal Reproduction, Ibaraki, Japan) were used. They were housed individually and kept on a 12 h light/dark cycle (lights on at 8:00 a.m.) and provided *ad libitum* access to food and water throughout the experiments. All experimental tests were conducted during the light phase. All experiments were approved by the University of Tsukuba Committee on Animal Research.

### Apparatus

Two open-field arenas (900 × 900 × 450 mm) made of black polyvinyl chloride or white acrylic plexiglass were used to test under two different contexts. The black context consisted of a gray floor and black walls. On one of the walls, a white–black vertically striped pattern was attached as a spatial cue. The white context consisted of a white floor and walls. A white–black checkered pattern was attached on one of the walls. The illumination at the center of each arena was 60 lx. An overhead camera was used to record the movement of the rat for the analysis. Background white noise (50 dB) was continuously present during all experimental tests to mask any extraneous noise. The stimulus objects were copies of 10 different objects made of glass, metal, or plastic and varied in height between 7 and 15 cm. All the objects were adequately heavy or fixed on the heavy metal plate such that the rat could not move them.

### Habituation

Habituation sessions were conducted for 3 consecutive days. On each day, rats received 5 min of handling by an experimenter, and were then placed in each of the black and white contexts without any objects for 30 min (with at least 60 min interval). The order of the exposure to each context was counterbalanced.

Following the habituation, rats were divided into three groups according to the length of the familiarization periods (5min, n = 11; 15min, n = 11; 30min, n = 10). All rats were subjected to three kinds of spontaneous recognition tests: OPC, place, and object recognition tests.

### Object-place-context recognition test

The OPC recognition test consisted of two sample phases and a test phase (Fig. 1A). A 24-hour delay period was inserted between the second sample and test phases. In the first sample phase (sample 1), rats were allowed to explore the black context where two different objects were diagonally placed at 22.5 cm apart from the adjacent two walls (e.g., object A on the top left and object B on the bottom right). In the second sample phase (sample 2), the rats were placed at the other white context in which the same pair of objects were placed in a swapped position relative to that in the first sample phase (e.g., object B on the left and object A on the right). Each rat was allowed to explore these objects freely for 5, 15, or 30 min at each sample phase. In the 5-min test phase, rats explored a pair of one of the sample objects (e.g. object A-A) in one of the contexts (e.g. black context). If rats had associative recognition memory of objects, places, and contexts, they would show a preferential exploratory behavior towards the object placed in the novel place-context combination (the dashed arrow in Fig. 1A). The positions of the novel objects (e.g. top left or bottom right) in the test phase were counterbalanced. After each phase, the floor of the arena was cleaned using a wet cloth containing sodium hypochlorite solution and the objects were wiped with 70% ethanol to eliminate odor.

**Fig. 1.**
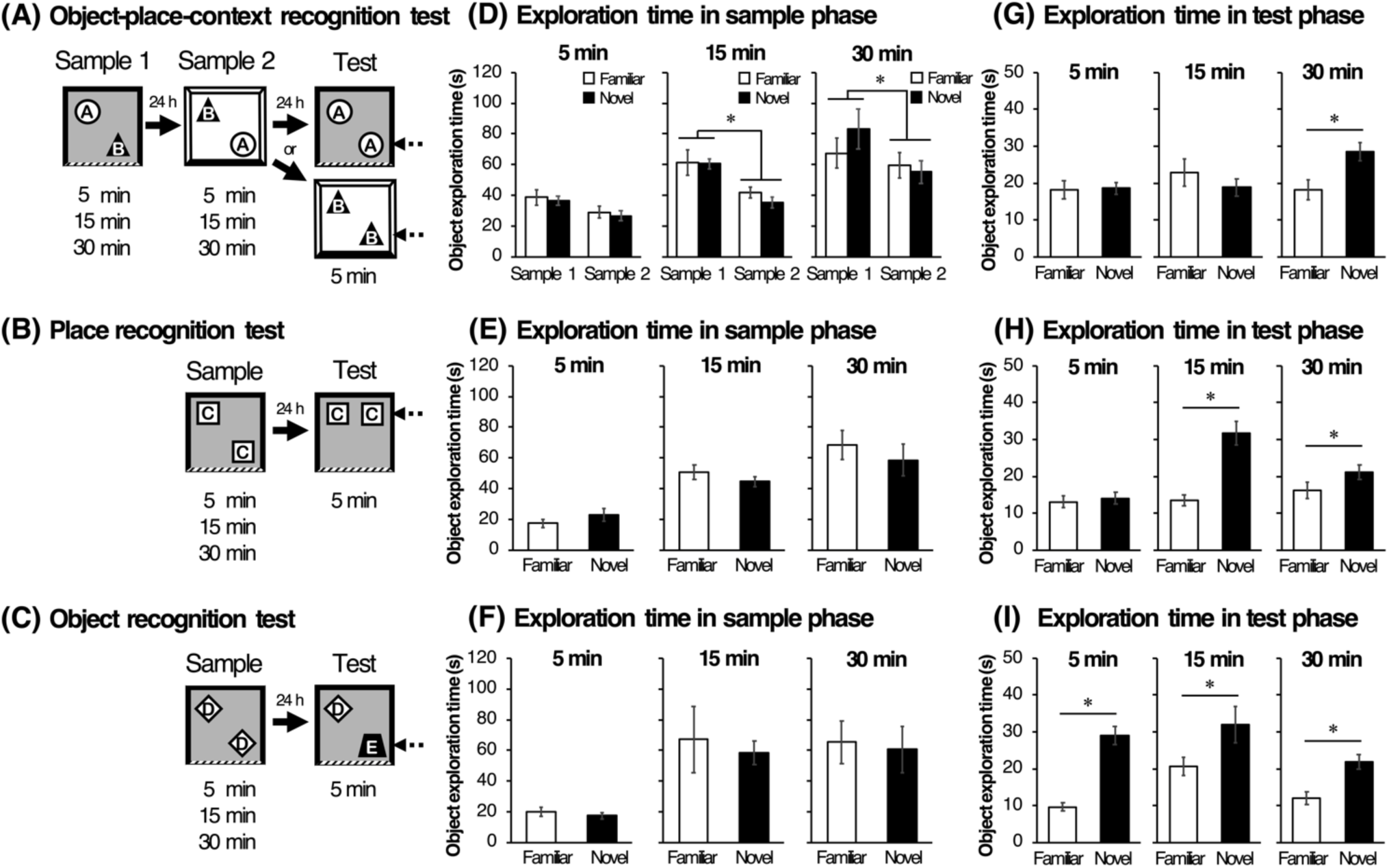
Schematic illustrations of object-place-context (OPC) recognition test (**A**), place recognition test (**B**), and object recognition test (**C**). Each test consists of sample phase (5, 15 or 30 min), delay interval (24 hours), and test phase (5 min). A successful discrimination is defined as a preferential exploration of the ‘object in a novel context-place combination’ in OPC recognition test, ‘the object in a novel location’ in place recognition test or ‘the novel object’ in object recognition test and is indicated by dashed arrows. Mean (±*SEM*) time spent in exploration for each object in sample phase of OPC recognition test (**D**), place recognition test (**E**), and object recognition test (**F**). Mean (±*SEM*) time spent in exploration for each object in test phase of OPC recognition test (**G**), place recognition test (**H**), and object recognition test (**I**).

### Place recognition test

Three-seven days after the OPC recognition test, the rats were subjected to a place recognition test. All rats were subjected to re-habituation to the black context for 15 min without any objects. Two identical objects (object C) and the black context were used in this test (Fig. 1B). Animals were allowed to explore two objects for 5, 15, or 30 min in the sample phase. After a 24-hour delay interval, the animals were returned to the context in which one of the objects were moved to a novel location and allowed to freely explore for 5 min in the test phase. Note that for the place recognition test, one of the objects was presented in the same location as familiar, whereas the other was moved to a different location (dashed arrow in Fig. 1B), which was placed 30 cm apart from the familiar object and 22.5 cm apart from a sidewall (two locations were possible).

### Object recognition test

3-7 days after the place recognition test, rats were tested in an object recognition test (Fig. 1C). All rats were subjected to re-habituation in the black context without the spatial cue for 15 min. In the object recognition test, the animals were allowed to explore two identical objects (object D) for 5, 15, or 30min in the sample phase. After the delay interval, animals were returned to the same open-field arena where one of the objects are replaced with a novel object (object E) in the 5 min test phase. The positions of the novel object in the test phase were counterbalanced.

### Data analysis

The ANY-maze video tracking software (Stoelting Co., Illinois, USA) was used to analyze the rats’ performance and locomotor activity in each test. In each phase, we manually counted the time rats spent exploring the objects. Exploration was defined as the rat sniffing, pawing, and directing its nose toward the objects within a distance of 2 cm, except standing over or climbing on the objects. As a measure of discrimination behavior in the test phase, discrimination index (DI) was calculated by dividing the difference in the time spent exploring the novel and familiar by the total time of exploration for both objects [DI = (T_novel_ − T_familiar_) / (T_novel_ + T_familiar_)]. A value of zero indicates no preference, while a positive value indicates more exploration of the novel object and a negative value indicates preferential exploration of the familiar object. Exploration time in the OPC, place recognition, object recognition test at each phase was analyzed using a paired Student’s t-test (two-tailed) or two-way (object × trial) analysis of variance (ANOVA) followed by a post-hoc Scheffe test. DIs were compared to the theoretical chance level (0%) using a one-sample t-test (two-tailed). An exclusion criterion of a statistical outlier was defined as occurring when DI exceeded ± 2 standard deviations from the mean of all rats in each test. If subjects met the criterion, the data in sample and test phases was excluded from the analysis. All values are expressed as mean ± standard error of the mean (SEM). Statistical significance was set at α= 0.05.

## Results

### Object-place-context test

In the 5 min condition, one subject was excluded from analyses according to the statistical outlier criterion. Fig. 1D shows the mean exploration time for the familiar and novel objects in the sample phases 1 and 2. Note that ‘familiar’ refers to the object that will be the same, and ‘novel’ refers to the object that will be replaced with a novel object in the test phase. A repeated two-way ANOVA, with Object (familiar vs. novel) as a within-subjects variable × Trial (sample 1 vs. sample 2) as a between-subjects variable, showed a significant main effect of Trial in the 30 min [F(1, 9) = 7.91, p = .020] and 15 min [F(1, 10) = 26.97, p = .004] conditions but not in the 5 min condition. A main effect of Object and an interaction between Object and Trial were not significant in each condition. Fig. 1G shows the mean exploration time for the familiar and novel objects in the test phase. Paired t-tests revealed that rats explored the novel object significantly more than familiar one (t(9) = 3.14, p = .011) in the 30 min condition. The mean DIs of each condition are shown in Fig. 2. One-sample t-tests revealed that DI was significantly higher than chance level only in the 30 min condition (t(9) = 2.85, p = .018).

**Fig. 2.**
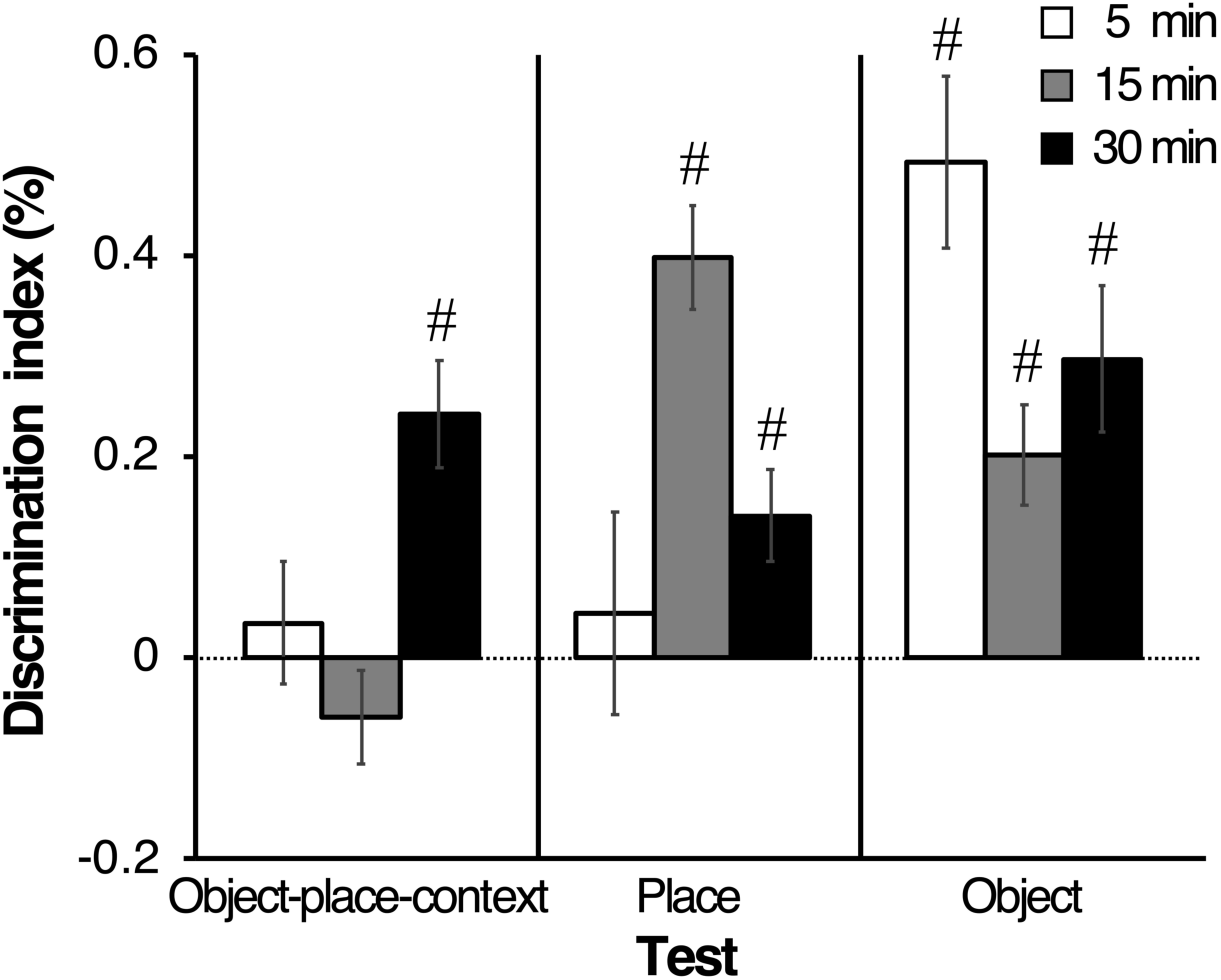
Mean (±*SEM*) discrimination index in test phase of OPC recognition test, place recognition test, and object recognition test. The horizontal dotted line indicates chance level (0). # *p*<.05 compared to chance level.

### Place recognition test

In the 5 min and 30 min conditions, one subject was excluded from analyses according to the statistical outlier criterion. Fig. 1E shows the mean exploration time for the familiar object and novel objects in the sample phase. A paired t-test revealed that there was no significant difference in the exploration time for each object in the sample phase. In the test phase, the mean exploration time for each object (Fig. 1H) and DIs in each familiarization condition (Fig. 2) were compared. A paired t-test revealed that rats explored the novel object significantly more than the familiar one when the sample phases were 15 min (t(10) = 5.73, p <.001) and 30 min (t(8) = 2.57, p = .032). Furthermore, one-sample t-tests revealed that DIs were significantly higher than chance level in the 15 min (t(10) = 7.66, p <.001) and 30 min (t(8) = 2.83, p = .022) conditions.

### Object recognition test

In the 15 min and 30 min conditions, one subject was excluded from analyses according to the statistical outlier criterion. Fig. 1F shows the mean exploration time for the familiar object and novel objects in the sample phase. A paired t-test revealed that there was no significant difference in the exploration time. In the test phase, the mean exploration time for each object (Fig. 1I) and DIs under three different familiarization conditions (Fig. 2) were compared. A paired t-test revealed that rats explored the novel object significantly more than the familiar one in all conditions (5 min, t(10) = 6.74, p <.001; 15 min, t(9) = 3.32, p =.008; 30 min, t(8) = 3.48, p =.008). One-sample t-tests also revealed that DIs were significantly higher than chance level in all conditions (5 min, t(10) = 9.19, p <.001; 15 min, t(9) = 4.39, p =.001; 30 min, t(8) = 3.87, p =.004).

## Discussion

In the present study, we investigated the relationship between the length of familiarization at the sample phase (5, 15, 30 min) and subsequent novelty discrimination performance at the test phase in the object, place and OPC recognition tests with a 24-hour delay period. In the OPC recognition test, rats showed a significant novelty preference when the familiarization period was 30 min, but not when it was 5 min or 15 min. In contrast, rats showed successful discrimination even under the shorter familiarization conditions such as 15 min in the place recognition and 5 min in the object recognition. These results demonstrated that the long-term (24 hours) associative recognition memory could be evident by extending the familiarization period to 30 min in the OPC recognition test. Furthermore, it is also suggested that the formation of the long-term complexed associative memory required longer familiarization compared to the simple non-associative memory.

The findings that successful recognition memory was evident in 5-min familiarization in the object recognition test, but not in the place recognition test, are consistent with our previous study showing that rats needed longer familiarization in the place recognition test than the object recognition test (Ozawa et al. 2011). Since rats are required to process the information on both objects and locations in the place recognition, it is reasonable that rats needed more time to process complexed information in the place recognition test than in the object recognition test. Thus, our results showing the relationship between the performance of recognition memory tests and the length of familiarization periods are likely to reflect the differences in difficulty among the object, place and OPC recognition tests. Previous studies reported that primates and humans spent more time gazing at a novel image than a familiar one in the habituation-dishabituation paradigm, and the longer familiarization period is required when the stimulus is more complex (Fagan 1964; Lasky 1980; Bachevalier et al. 1993). Although Gaskin et al. (2010) demonstrated that a longer exploration of the objects in the familiarization period did not improve non-associative memory performance in the object recognition test of rodents, our findings, which showed that the longer the familiarization period, the more time rats spent in exploration for the objects in the sample phase, demonstrated that the formation of associative recognition memory needs much longer exploration for the objects. These results suggest that a sufficient exploration of the environment, as well as objects and/or locations, can lead animals to make associations between each elements of information, such as objects, locations, and contexts.

Associative recognition memory has been regarded as an episodic-like memory, which is typified as the comprehensive information of “what”, “where” and “when” acquired from “a single experience” (Eacott and Norman 2004). Indeed, although several tests have been developed to measure episodic-like memory in rodents, some of them are thought to be inappropriate for episodic-like memory tests. For example, a food reinforcement-based test (Veyrac et al. 2015) is unlikely to meet the definition of episodic-like memory because it requires multiple training sessions. According to its definition, the test for episodic-like memory should be completed in a few training sessions. In addition, direct comparison between associative and non-associative memories is thought to be impossible, even in reinforcement-free spontaneous recognition tests, due to differences in the procedures of the tests. The associative memory test (episodic-like memory test: Dere et al. 2005; Kart-Teke et al. 2006) used more objects (e.g. 4 objects that have different memory properties including the object, place, and temporal element) in a single-trial test compared to the non-associative memory tests in which two objects are usually used. In other words, the performance in the episodic-like memory test using multiple objects in a single trial could depend on how much animals focus on each memory element (object, place, and temporal) in the test phase. In the OPC recognition test, however, only two objects are used and multiple training is not required. Also, it includes the elements of episodic-like memory. Thus, the OPC recognition test can be the most appropriate paradigm for a rodents’ episodic-like memory test, and it can systematically investigate the cognitive and neural mechanisms in both associative and non-associative memory by combining with object recognition and place recognition tests.

In conclusion, our results showed that long-term associative recognition memory was evident when the familiarization period was extended to 30 min in the OPC recognition test. The findings suggested that longer familiarization periods are necessary for the recognition of the complex associative memory compared to simple non-associative memory. We propose that a spontaneous recognition paradigm is a useful tool for the systematic assessment of long-term associative and non-associative recognition memory in rats.

## Acknowledgements

This study was supported by JSPS KAKENHI Grant Numbers 18H01097, JP19K03365, JP19K21806.

